# Attentional modulation of the cortical contribution to the frequency-following response evoked by continuous speech

**DOI:** 10.1101/2023.07.03.547608

**Authors:** Alina Schüller, Achim Schilling, Patrick Krauss, Stefan Rampp, Tobias Reichenbach

## Abstract

Selective attention to one of several competing speakers is required for comprehending a target speaker amongst other voices and for successful communication with them. Selective attention has been found to involve the neural tracking of low-frequency speech rhythms in the auditory cortex. Effects of selective attention have also been found in subcortical neural activities, in particular regarding the high-frequency neural response at the fundamental frequency of speech, the speech-FFR. Recent investigations have, however, shown that the speech-FFR contains cortical contributions as well. It remains unclear whether these are also modulated by selective attention. Here we employed magnetoencephalography (MEG) to assess the attentional modulation of the cortical contributions to the speech-FFR. We presented participants with two competing speech signals and analyzed the cortical responses during attentional switching between the two speakers. Our findings revealed robust attentional modulation of the cortical contribution to the speech-FFR: the neural responses were higher when the speaker was attended than when they were ignored. We also found that, regardless of attention, a voice with a lower fundamental frequency elicited a larger cortical contribution to the speech-FFR than a voice with a higher fundamental frequency. Our results show that the attentional modulation of the speech-FFR does not only occur subcortically but extends to the auditory cortex as well.

## Introduction

Selective attention is a fundamental cognitive process that allows us to focus on relevant information while filtering out distracting signals. Referred to as cocktail-party effect, in complex acoustic settings such as a busy pub or restaurant, selective attention enables us to, for instance, focus on a particular speaker amongst other competing voices to selectively process that speech signal and extract linguistic information and meaning (1; 2).

Recent research has employed continuous natural speech to explore which speech features are involved in selective attention to speech. These investigations have mostly focused on low-frequency responses in the auditory cortex. In particular, they found that the neural tracking of rhythms in speech set by the rate of syllables and words, in the delta (1–4 Hz) and theta (4–8 Hz) frequency bands, is modulated by selective attention to one of two competing speakers (3; 4; 5; 6; 7). In addition, the power of high-frequency responses in the gamma band, between 70 - 150 Hz, tracks the low-frequency speech rhythms as well (8; 9). These findings were obtained using different measurement techniques, in particular an invasive one, electrocorticography (ECoG) (8), as well as non-invasive ones such as magnetoencephalography (MEG) (4) and electroencephalography (EEG) (3; 5; 10).

As evidence of the robustness of the attentional effect on the neural tracking, attention to a specific voice could be accurately decoded from single trials with short speech stimuli that lasted approximately one minute using MEG (4) and EEG (11; 12; 13). The decoding accuracy further improved with optimization of statistical modeling, allowing for accurate decoding from recordings shorter than 30 seconds (14; 15). Additionally, the ability to detect changes in attentional focus within tens of seconds was demonstrated using EEG data, and even faster when combined with sparse statistical modeling techniques in MEG data, as observed in the study by (16).

In addition to the cortical tracking of the low-frequency speech rhythms, a high-frequency neural response to a high-frequency speech features has been investigated as well (17; 18). It emerges in response to the fundamental frequency and its higher harmonics of the voiced parts of speech. The frequency range of the response is that of the fundamental frequency of speech, typically between 100 Hz and 300 Hz. Because of its similarities to the frequency-following response (FFR) to a pure tone, we refer to it as the *speech-FFR* in the following.

Using EEG, we have recently shown that the speech-FFR is modulated by selective attention to one of two competing speakers (19; 20; 21). In particular, we measured a larger speech-FFR to a particular speaker when that speaker was attended as compared to when they were ignored. The latency of the speech-FFR that we analyzed was about 10 ms. Together with the topographic map that showed large contributions from the mastoid channels and the channels at the vertex, this demonstrated a subcortical origin of the response. The attention decoding accuracy was significant even for short segments of EEG data, down to a few seconds in duration (20).

However, studies using MEG recently showed the presence of cortical contributions to the speech-FFR (22; 23; 24; 25; 26; 27). These contributions have been verified through EEG measurements as well (28). In response to continuous speech, the cortical portion of the speech-FFR has been found to occur at latencies of around 30-40 ms (26; 29). EEG and MEG have thereby been found to play partly complimentary roles when assessing the speech-FFR: EEG measures mostly the subcortical contributions, while MEG records predominantly the cortical portions.

Whether the cortical contribution to the speech-FFR is modulated by selective attention has, however, not yet been investigated. Here we set out to close that gap in knowledge. In particular, by employing the sensitivity of MEG to cortical sources as well as an auditory stimulus that contained two competing continuous speech signals, we aimed to investigate how attention modulates cortical responses during continuous speech perception.

## Materials and Methods

### Experimental design and data analysis

We employed MEG recordings of neural responses to two competing talkers (Figure 1a). The speech signals consisted of two audiobooks, and participants regularly switched their attention between the two speakers. The MEG data was analyzed through first performing source reconstruction and then relating the source-reconstructed neural activity to two high-frequency speech features using linear regression. We thus obtained temporal response functions (TRFs) that described the speech-FFR. We then compared the TRFs between the condition in which the corresponding speaker was attended to the condition where they were ignored.

**Figure 1:**
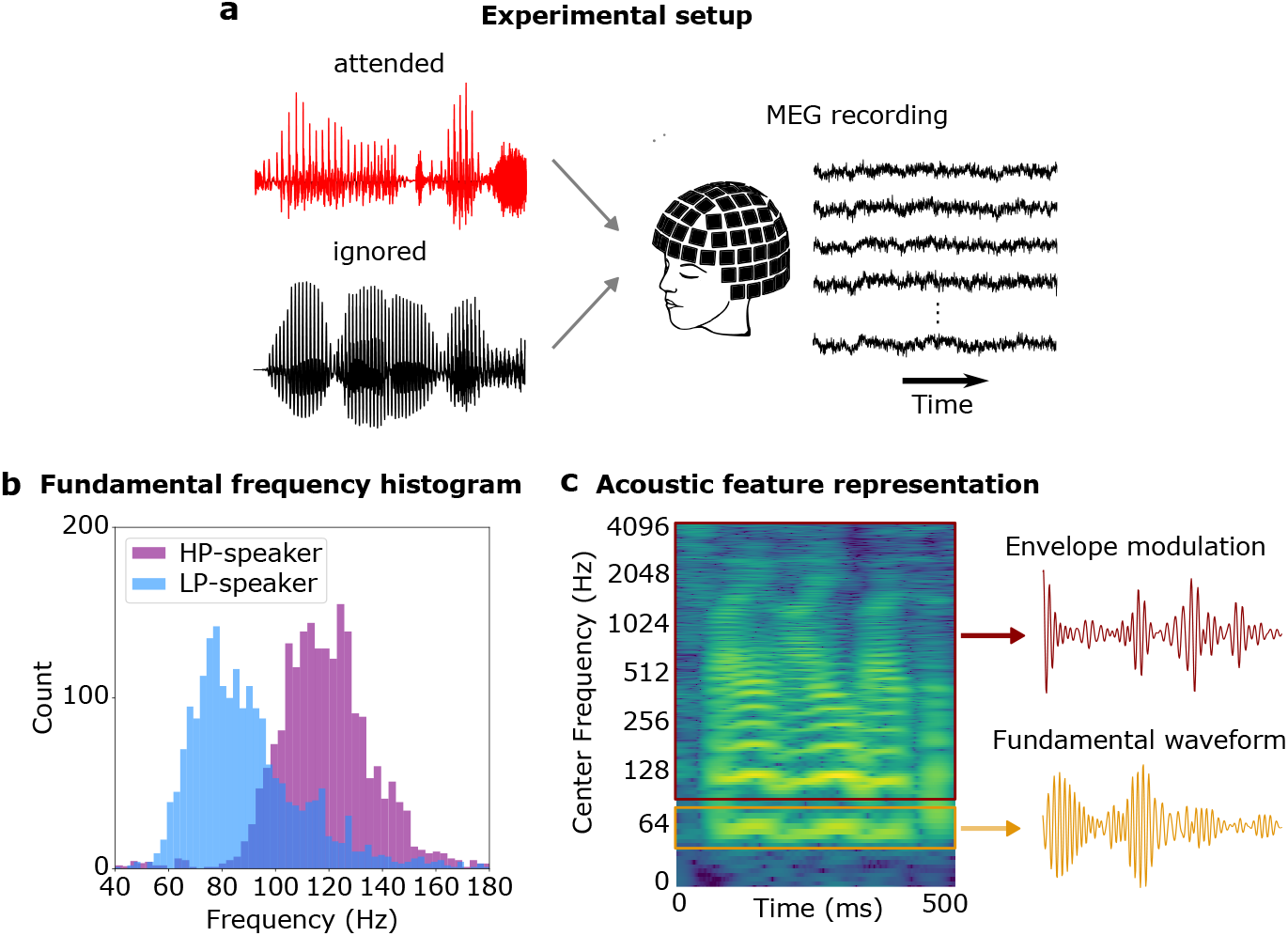
Experimental setup and acoustic stimuli. (a) Two audiobooks (one attended and one ignored) were presented simultaneously while MEG was recorded. (b) One of the two male speakers had a lower fundamental frequency and hence pitch (LP) and the other speaker a higher one (HP). (c) We quantified the speech-FFR through two acoustic features: the *fundamental waveform* that reflected the portion of the speech spectrogram around the fundamental frequency, and the *envelope modulation* of the higher harmonics.

### Participants

We recruited 22 healthy, right-handed, native German speakers (10 females, 12 males, 19-29 years) with no history of neurological disease or hearing impairment. The study was granted ethical permission by the ethics board of the University Hospital Erlangen (registration no. 22-361-S).

### Speech stimuli

The participants listened to approximately 40 minutes of acoustic stimuli consisting of two German audio-books (referred to as *story-audiobooks*) narrated by two competing male speakers. Two additional German audiobooks (referred to as *noise-audiobooks*) of the same narrators were used to serve as background noise.

The first story-audiobook was “*Frau Ella*” by Florian Beckerhoff, and the second story-audiobook was “*Den Hund überleben*” by Stefan Hornbach. As the first noise-audiobooks we employed “*Darum*” by Daniel Glattauer, and as the second noise-audiobook “*Looking for hope*” by Colleen Hoover (translated to German by Katarina Ganslandt). The first story-audiobook and the first noise-audiobook were narrated by Peter Jordan, and the second story-audiobook and the second noise-audiobook were narrated by Pascal Houdus. All audiobooks were published in *Hörbuch Hamburg* and are available in stores.

Peter Jordan’s voice had, on average, a lower pitch than the voice of Pascal Houdus (Fig. 1b). In particular, the fundamental frequency of Peter Jordan’s voice varied between 70 - 120 Hz, whereas that for Pascal Houdus occured between 100 - 150 Hz. We therefore refer to Peter Jordan as the *lower-pitch (LP) speaker* in the following, and to Pascal Houdus as the *higher-pitch (HP) speaker*.

The stimuli containing the two competing speakers were presented diotically, at approximately the same sound-pressure level of 67 dB(A) throughout the experiment. In order to facilitate the listening process for the participants, we kept the original chapters of the story-audiobooks and divided them into segments of lengths according to the original chapter lengths. The resulting chapter lengths were between 3 and 5 minutes long. The background noise was generated by randomly picking audio segments from the noise-audiobooks that where of the same length as the chapters from the story-audiobook. In total we employed approximately 37 min of acoustic stimuli.

### MEG data acquisition

Two speech stimuli were presented simultaneously, with the first chapter of the first story-audiobook played alongside an unrelated story narrated by the speaker of the second noise-audiobook in the background. Subsequently, the first chapter of the second story-audiobook was played alongside an unrelated story narrated by the speaker of the first noise-audiobook in the background. This pattern continued so that both story-audiobooks were told in a subsequent but alternating manner.

To assess the impact of selective attention on the cortical response, the participants were instructed to selectively attend only one of the two acoustic stimuli, the story-audiobook (Fig. 1a. In detail, the participants switched their attention between the two speakers with every chapter, starting by attending the LP speaker for the first chapter of the first audiobook, then attending the HP speaker in the following, for the first chapter of the second audiobook, then again attending the LP speaker for the second chapter of the first audiobook, and so on. In order to instruct the participants which speaker they should attend, the story-audiobook always started alone and about 5 seconds later the noise-audiobook started. After each chapter, before attention switched, participants were visually presented with three single-choice questions to assess whether they correctly attended the intended speaker. The total stimulation protocol lasted approximately 50 to 55 minutes.

MEG data was recorded using a 248 magnetometer system (4D Neuroimaging, San Diego, CA, USA) with a sampling frequency of 1,017.25 Hz. The participants were in a supine position with their eyes open during the recordings. An analogue bandpass filter (1.0 − 200 Hz) was applied online to eliminate unwanted frequency components. To correct for environmental noise, a calibrated linear weighting of 23 reference sensors (manufacturing algorithm, 4D Neuroimaging, San Diego, CA, USA) was applied and five landmark positions were recorded using an integrated digitizer (Polhemus, Colchester, Vermont, Canada). Additionally, prior to each measurement, head shape digitization was performed. For further analysis, a 50 Hz notch filter (firwin, transition bandwidth 0.5 Hz) was applied offline using MNE-Python (30), to remove power line interference, the data was downsampled to a sampling frequency of 1,000 Hz to facilitate subsequent processing and underwent offline digital band-pass filtering (between 70 - 120 Hz for the LP voice, between 100 - 150 Hz for the HP voice; linear digital Butterworth filter, second order, critical frequencies obtained by dividing the lower and upper cut-off frequency by the Nyquist frequency, applied forward and backward).

The speech signal was presented during the MEG recordings using a custom-designed setup, as described in detail in our previous study on linguistic responses (31) and in our previous study on the speech-FFR (29). The setup involved a stimulation computer connected to an external USB sound device with five analog outputs. Two of these outputs were connected to an audio amplifier. The first output was linked in parallel to an analog input channel of the MEG data logger, recording the mixed audio stimulus as presented to the subject. To achieve precise alignment between the speech stimulus and the MEG recording, we employed cross correlation of the speech stimulus with the audio reference recording obtained from the analog input channel of the MEG data logger, yielding an alignment accuracy of 1 ms.

### Acoustic stimulus representations

We employed two acoustic speech features to capture the high-frequency neural response at the fundamental frequency of the speech signals: the fundamental waveform *f*_0_(*t*) and the higher-mode envelope modulation *e*(*t*) (Fig. 1c). The fundamental waveform was extracted using probabilistic YIN (pYIN) algorithm (32). As an adaptation of the YIN algorithm (33), pYIN is a method employed for the estimation of the fundamental frequency (*f*_0_). By integrating the YIN algorithm and Viterbi decoding (34), pYIN enhances the accuracy and reliability of the *f*_0_ estimation. The utilization of YIN initially generates a set of potential *f*_0_ candidates, whereas the subsequent application of Viterbi decoding refines these candidates, producing a more precise estimation of the *f*_0_ contour. This combined approach provides an effective methodology for extracting fundamental frequency information from audio signals.

We extracted the fundamental frequency *f*_0_ for each chapter of the LP audiobook separately, resulting in a cutoff-frequency for the *f*_0_-filtering of 65 Hz for the lower edge and of 120 Hz for the upper edge. For the HP audiobook we did the same, yielding a cutoff-frequency of 99 Hz for the lower edge and of 145 Hz for the upper edge. These estimated frequency bands matched well with the fundamental frequency histograms obtained for both speakers (Fig. 1b.

As already demonstrated in previous studies (35; 26; 29), the envelope modulation of the higher harmonics in an acoustic signal contributes to the neural response at the fundamental frequency even more than the fundamental frequency itself. To describe these envelope modulations, we employed a computational model of the auditory periphery that draws inspiration from both psychoacoustics and neurophysiology. It aims to replicate and simulate the fundamental processes and mechanisms underlying auditory perception by taking into account the tonotopic organization of the cochlea, the frequency tuning properties of auditory-nerve fibers, and the neural responses observed in various auditory nuclei along the ascending pathway. The model was originally implemented within the NSL auditory-cortical toolbox in Matlab (36). For this study, we used the parts of this toolbox that describe the early auditory processing. The Matlab code was translated to Python within our group.

In detail, the model employs firstly a bank of constant-*Q* bandpass filters, that is, specialized filters that have a varying frequency resolution across the auditory spectrum. These filters are followed by nonlinear compression and derivative mechanisms implemented across different scales, yielding a sharpening of frequency resolution. An envelope detector is then applied to each frequency band that extracts the amplitude envelope of the signal at each specific frequency band.

### Neural source estimation

The source reconstruction process was implemented using the MNE-Python software package (30). Given the absence of subject-specific MR scans, we employed the Freesurfer template MRI called ‘fsaverage’ as a substitute (37). Previous studies have shown that using an average brain template can yield results comparable to individual MR scans in source localization analyses (38; 39). Its validity has not only been established in our recent study on early subcortical MEG responses to continuous speech (29), but also in previous studies that investigated high-frequency neural mechanisms of speech processing (26).

To account for individual differences, we collected information about the head position of each subject with respect to the MEG scanner at the beginning and end of each measurement, using five marker coils. Additionally, we digitized the shape of each subject’s head using a Polhemus system (Colchester, Vermont, Canada). The so-obtained subject-specific information was then employed to align the ‘fsaverage’ brain template with each individual’s head by applying rotation, translation, and uniform scaling.

To create a volumetric source space for the average brain, we employed a regular grid with neighboring grid points spaced at intervals of 5 mm. This volume source space was subjected to the application of the Freesurfer ‘aparc+aseg’ parcellation, to subsequently define a specific region of interest (ROI) for source estimation.

We established a cortical ROI, representing the auditory cortex in both the right and left hemispheres. Specifically, the ‘aparc’ labels ‘transversetemporal’, ‘middletemporal’, ‘superiortemporal’, ‘supramarginal’, ‘insula’, and ‘bankssts’ were employed to identify the cortical ROI. This cortical subdivision resulted in a total of 525 source locations with arbitrary orientations.

In order to generate a volume conductor model that accurately represents the shape of the head for source reconstruction, even in the absence of subject-specific MR scans, we employed the boundary element model provided by Freesurfer for the fsaverage brain template. Using the volume source space and the forward solution, we computed a linearly constrained minimum variance (LCMV) beamformer (40). The LCMV beamformer is a spatial filter that applies a set of weights to scan through each source location in the predefined source space grid. It independently estimated the MEG activity at each source point.

To perform the source reconstruction, we used a data covariance matrix estimated from a one-minute segment of MEG data acquired during audio stimulation, as well as a noise covariance matrix derived from three-minute pre-stimulus empty room recordings. The beamformer filter was then applied to the raw but pre-processed MEG data of each subject. This process resulted in the estimation of a three-dimensional current dipole vector, with magnitude and direction, at each source location. We thus source-reconstructed the pre-processed MEG data of every subject.

### Temporal Response Functions (TRFs)

To investigate the latency and origin of the measured neural signal, we employed a linear forward model that aimed to predict the neural activity 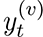 of each source point (voxel) *v* at time *t* from a linear combination of the acoustic stimuli *f_t_* and *e_t_*. The acoustic stimuli were shifted by different time delays *τ*. The resulting weights 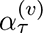 and 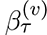 of this linear equation are the temporal response functions (TRFs), one for each source point. A TRF provides a quantitative representation of how the system’s output changes over time due to changes in the input. Thus, the TRF for each voxel describes the corresponding neural response to each acoustic feature across a range of time delays, from *τ*_min_ to *τ*_max_:

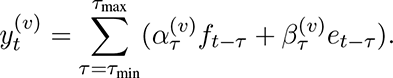

We used regularized ridge regression to estimate the TRFs. In this approach, the regularization parameter *λ* can be defined as *λ* = *λ_n_ · e_m_*, where *λ_n_* represents the normalized regularization parameter, and *e_m_* corresponds to the mean eigenvalue of the covariance matrix. After applying five-fold crossvalidation to estimate the appropriate regularization parameter for each subject, we found that the same regularization parameter *λ* = 1 could be used for all subjects, since the highest model accuracy appeared close to this value for all subjects. The forward model and the TRF estimation used in this study were implemented in Python based on the algorithms developed previously in our group by (20) and (35).

We considered a range of time delays of *τ*_min_ = −20 ms to *τ*_max_ = 120 ms with an increment of 1 ms since the sampling rate was 1,000 Hz. This resulted in 141 time lags in total. We calculated both the voxel-wise TRFs on the subject-level to capture subject-specific diversity, as well as TRFs on the population average to provide representative results as well as statistical inference on population level.

### Attentional modulation of the cortical response

In order to investigate the impact of attention on the cortical contribution to the speech-FFR, we computed two pairs of TRFs for each acoustic feature and each subject.

The first pair of TRFs respresented the neural response to the LP speaker. One TRF was estimated when the LP voice was attended by the subject (referred to as the LP-A condition), while the other TRF was constructed when the LP was ignored (referred to as the LP-I condition).

Similarly, the second pair of TRFs was designed to represent the neural response to the HP speaker. The first TRF in this pair was computed when the subject directed their attention to the HP voice, referred to as the HF-A condition. The second TRF in the pair was created when the subject ignored the HP speaker, referred to as the HF-I condition,.

Previous studies found that the cortical portion of the speech-FFR, as assessed through TRF amplitude, occurs predominantly at delays between 30-40 ms (26; 23; 29). The TRF amplitudes are modulated by the fundamental frequency multiplied by two, since maxima and minima in the TRFs are both mapped to maxima when computing the amplitude. Several maxima and minima therefore arise in the TRF amplitude between 30-40 ms. Since the latencies between subjects may slightly vary in a range of a few ms, it is likely that at a time lag of 34 ms, for instance, one subject might display a maximum in the TRF amplitude and another subject a minimum. A direct comparison between the amplitudes of the TRFs in the attended and ignored conditions at one specific time lag may hence lead to inaccurate results. To avoid this problem, we computed the envelopes of the TRF amplitudes. The envelope represents the magnitude of the signal over time, capturing the overall modulation pattern without relying on precise temporal alignment. Using envelope comparisons allowed us to capture the essential characteristics of the cortical response to different speakers while minimizing the impact of small temporal variations. This approach provided a more stable and interpretable basis for evaluating the attentional effects on the cortical response.

### Significance of the cortical response

To assess the statistical significance of the neural responses at the population level, we conducted statistical tests by comparing the calculated TRFs to noise models. The noise models were generated by circularly shifting the audio feature in time, as described in (26) and already employed in (29).

Specifically, we performed circular shifts of each audio feature with time shifts of 15 seconds, 30 seconds, and 45 seconds, respectively. For each shifted audio feature, we computed the corresponding TRFs and then averaged the TRFs obtained from all time-shifted features. This process was repeated for each subject, resulting in two noise models (one for each audio feature) per subject.

To evaluate the statistical significance of the TRFs, we employed a bootstrapping approach with the single-subject noise models for each audio feature. This process involved resampling the noise models across time lags and subjects, averaging them across subjects and vertices, and computing magnitudes over time lags in a manner consistent with the actual TRFs. By repeating this procedure 10,000 times, we generated a distribution of noise model magnitudes across time lags. We then determined the proportion of values from this noise distribution that exceeded the magnitude of the actual TRF for each model. This allowed us to estimate empirical p-values for each time lag. To account for multiple comparisons, the estimated p-values were corrected using the Bonferroni method.

### Lateralization of the cortical responses

To investigate the potential lateralization of cortical activity at time delays where significant responses emerged, we conducted a two-tailed Wilcoxon signed rank test. This test focused on assessing the differences in magnitudes of the population-average TRFs between the right and left cortical ROIs at the time delays of significant responses.

### Significance of attentional modulation

To assess potential significant differences in the envelopes of the TRF magnitudes between the attended and the ignored condition for individual subjects, we conducted a two-tailed Whitney-Mann rank test at the subject level. We therefore split the source-reconstructed MEG data as well as the acoustic stimuli into ten segments of equal length and calculated for each of them the TRF amplitude and the corresponding envelope. We then extracted the envelope value of each split-TRF at a certain latency time, which was chosen based on the peak of the population-average TRF magnitude. We thus generated a distribution of magnitude values for each subject and each acoustic feature, which was then used for the statistical testing.

We additionally employed a two-tailed Wilcoxon signed rank test to investigate whether the population-average envelope of the TRF magnitude of each feature deviated significantly between the attended and the ignored condition.

Finally, we employed a two-tailed Wilcoxon signed rank test to examine whether the population-average TRFs obtained for the LP speaker and the HP speaker differed significantly.

## Results

### Evaluation of comprehension questions

To test whether the participants attended the target speaker, we presented three single-choice questions (in total 15 questions per audiobook narrator) at the end of each chapter. Each question had four response options, resulting in a 25% chance level. Regarding the LP speaker, the questions were answered with an accuracy of 81 *±* 3% (mean and standard error of the mean). The response accuracy for questions related to the HP speaker was comparable at 80 *±* 2%.

### Cortical responses to the LP and the HP speaker

We measured neural responses to two competing male speakers, with distinct, but partly overlapping fundamental frequencies (Fig. 1b). The participants were asked to switch the attention from the LP to the HP speaker and back after each chapter of the corresponding audiobook. The recorded MEG data were then analyzed separately, first by source-reconstructing the preprocessed MEG signals and subsequently calculating source-level TRFs in a cortical ROI.

We then computed four linear forward models, each of which contained two acoustic features, the fundamental waveform and the envelope modulation. The model for the first condition, LP-A, captured the neural responses when the lower-pitch speaker was attended, while the model for the second condition, LP-I, represented the response to the ignored lower-pitch speaker. The models for the third and fourth condition, HP-A and HP-I, were calculated analogously to describe the neural responses when the HP speaker was attended or ignored, respectively.

We first verified that, for both acoustic features, the population-average TRFs showed significant neural activity in the auditory cortex. For the fundamental waveform, the TRF showed significant activity between 21 ms to 61 ms, peaking at 35 ms (*p <* 0.0001), when the LP speaker was attended and between 26 ms to 50 ms, peaking at 33 ms (*p <* 0.0001), when the LP voice was ignored (Fig. 2, upper left). The source activation at the peak time lags in both the LP-A condition and the LP-I condition showed a right lateralization (LP-A: *p* = 5.5 *·* 10*^−^*^12^; LP-I: *p* = 1.2 *·* 10*^−^*^7^).

**Figure 2:**
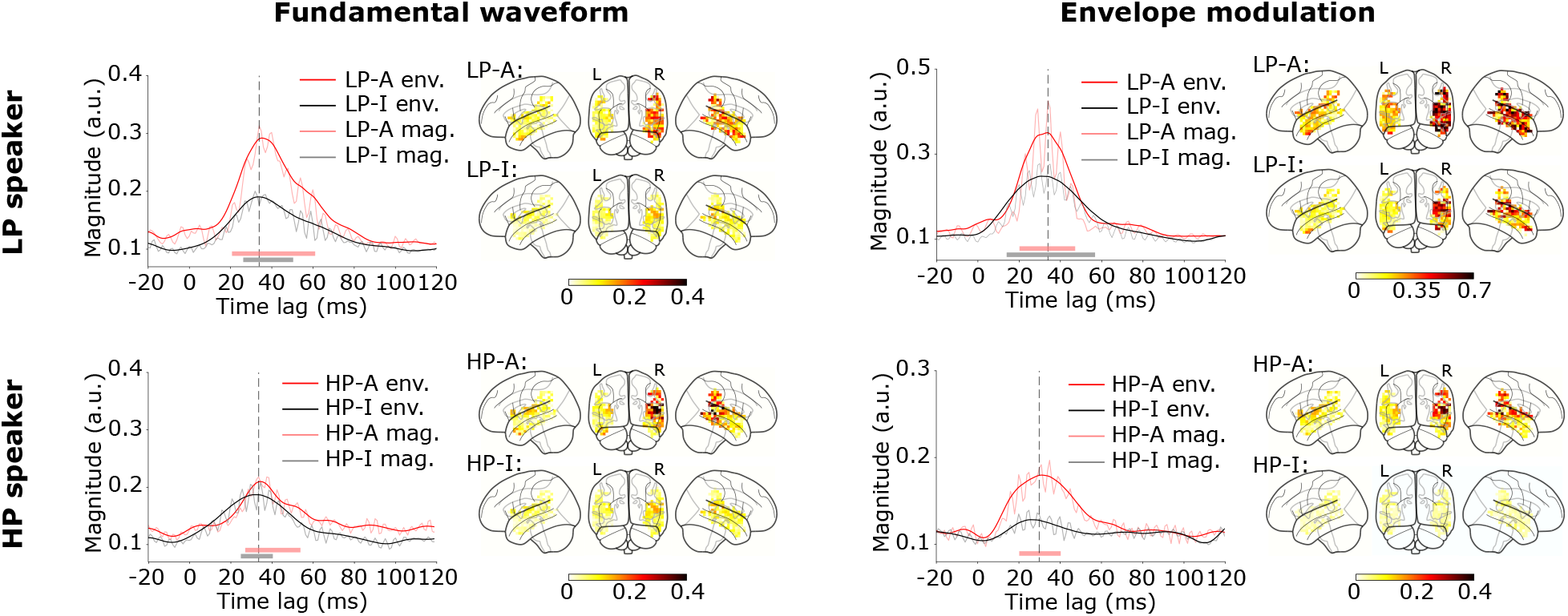
Cortical responses to both acoustic features of the LP and the HP voice. The voxel- and subject-averaged envelopes of the TRF amplitudes are significant at time lags around 34 ms (dashed grey lines) for both features and both in the LP-A condition (LP-A env., red) and in the LP-I condition (LP-I env., black). The TRF magnitudes display oscillation at 2*f*_0_ and are displayed as well (LP-A mag., pink; LP-I mag., grey). Significant time delays are indicated through the colored bars at the bottom of the plots. The corresponding brain plots show right-lateralized activity at 34 ms (attended, upper row; ignored, lower row). The same applies for the response to the fundamental waveform in the HP-A and in the HP-I condition. The envelope modulation for the HP voice shows significant activity at time lags around 30 ms (dashed grey line) when the speaker was attended, but not when he was ignored.

For the envelope modulation, the population-average TRFs for the LP-A condition yielded significant cortical activity at time lags between 20 ms and 47 ms, peaking at 34 ms (*p <* 0.0001). Regarding the LP-I condition, the TRFs showed significant activity at delays ranging from 14 ms to 57 ms, peaking at 31 ms (*p <* 0.0001; Fig. 2, upper right). As for the fundamental waveform, the source activation also revealed a right-lateralized dominance at the peak time lags in both the LP-A and the LP-I condition (LP-A: *p* = 1.5 *·* 10*^−^*^8^; LP-I: *p* = 2.3 *·* 10*^−^*^13^).

For the HP voice, the envelopes of the TRF amplitudes showed a similar behaviour. When attended, the TRF for the fundamental waveform yielded a significant response between 25 ms and 54 ms, peaking at 35 ms (*p <* 0.0001). The TRF in the HP-I condition showed significance between 25 ms and 40 ms, also peaking at 36 ms (*p <* 0.0001; Fig. 2, lower left). The source activation emerged similarly to the LP voice, and was right-lateralized at the peak time lags (HP-A: *p* = 5.2 *·* 10*^−^*^8^; HP-I: *p* = 6.4 *·* 10*^−^*^5^).

For the envelope modulation of the HP speaker, a significant response only emerged when the HP voice was attended, in a range from 20 ms to 40 ms, peaking at 36 ms (*p* = 0.02; Fig. 2, lower right). The TRF in the HP-I condition did not show any significant time lags. The source activation in the HP-A condition at the peak time lags exhibited a right-lateralization (HP-A: *p* = 1.9 *·* 10*^−^*^6^).

Neural responses for individual subjects in each of the four conditions LP-A, LP-I, HP-A, and HP-I will be presented below.

### Differences between responses to the LP voice and the HP voice

We wondered if the neural responses to the two speakers differed, independent of any putative attentional modulation. We hence investigated if the neural response to the LP speaker was systematically higher or lower than that to the HP speaker.

To assess those differences in the responses to the LP compared to the HP voice for each acoustic feature, we extracted from each subject the envelope of the TRF magnitude at the latency of interest (Fig. 2, dashed black lines). We then applied a two-tailed Wilcoxon signed rank test to assess whether there was a significant difference in the TRF amplitudes between the LP and the HP speaker. We did this separately for the attended and the ignored conditions.

Regarding the response to the fundamental waveform, we found a significantly higher envelope of the TRF amplitude for the LP speaker than for the HP speaker when the respective speaker was attended (*p* = 0.0049) (Fig. 3, upper left), but not when it was ignored (*p* = 0.58) (Fig. 3, lower left).

**Figure 3:**
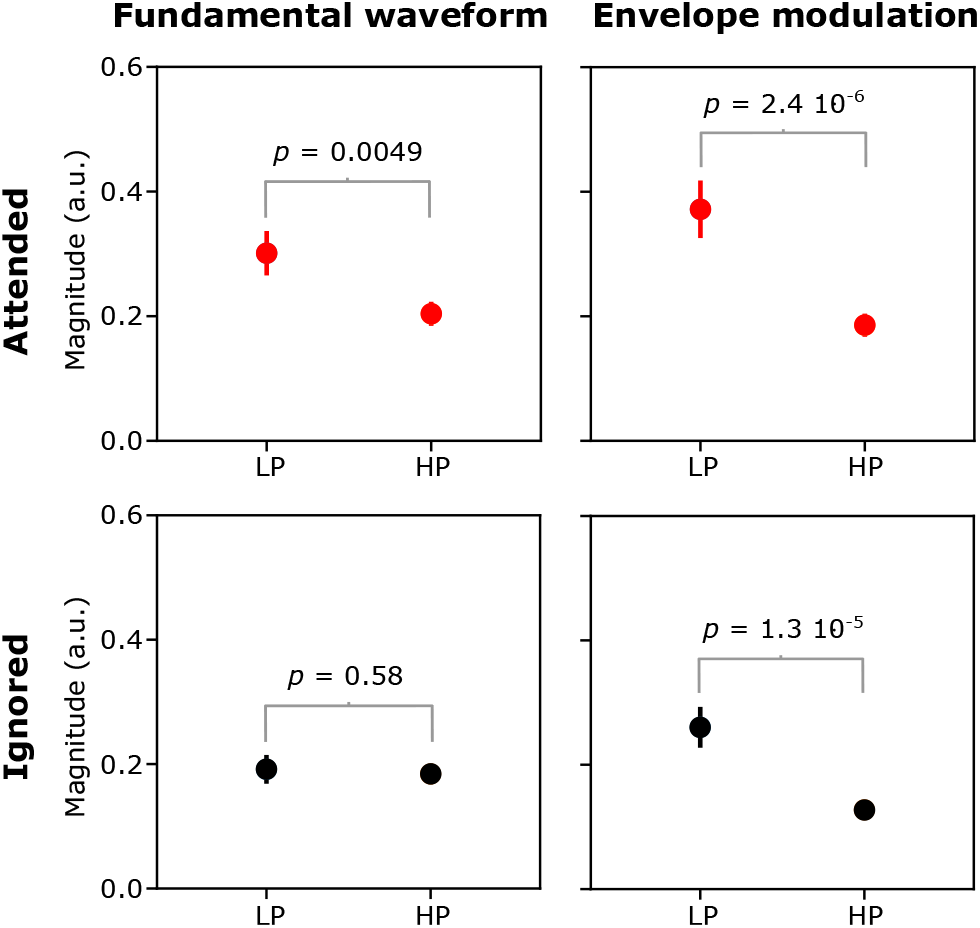
Comparison of the responses to the LP voice to the responses to the HP speaker. We show the peak of the envelope of the TRF magnitudes. The responses to the fundamental waveform (upper left) and to the envelope modulation (upper right) were significantly higher for the LP voice than for the HP voice when these voices were attended (two-tailed Wilcoxon signed rank test). When the voices were ignored, the difference between the responses to the LP voice and the HP voice were significant for the envelope modulation (lower left), but not for the fundamental waveform (lower right, two-tailed Wilcoxon signed rank test). The error bars indicate the standard error of the mean when averaging across subjects.

For the response to the envelope modulation, the comparison yielded a significant difference when both speakers were attended (*p* = 2.4 *·* 10*^−^*^6^) (Fig. 3, upper right), as well as when both speakers were ignored (*p* = 1.3 *·* 10*^−^*^5^; Fig. 3, lower right). In each case, the neural response to the LP voice was higher than that to the HP one.

### Attentional modulation of the cortical contribution to the speech-FFR: response to the LP speaker

To investigate how attention modulates the cortical response, we employed a classic paradigm of auditory selective attention paradigm, where participants attended one of two competing speech signals while ignoring the other one. We then assessed the neural responses to both speakers, and compared the responses to the same speaker when they were attended to the condition in which they were ignored.

To quantify the neural responses, we extracted the peak value of each of the four population-averaged envelopes of the TRF magnitudes (Fig. 2, dashed grey lines). For statistical analysis, we moreover calculated ten split-TRFs for each subject and each condition (LP-A, LP-I, HP-A and HP-I), extracted the value of the corresponding envelope of the TRF magnitudes at the prior extracted latency time of interest, and applied a two-tailed Whitney-Mann rank test on the difference between the values obtained for the attended condition and the ignored condition. This was done for the responses to the LP voice and the HP voice separately. It is important to note here that the following Figs. 4, 5, 6 and 7a all show the values of the attended and ignored TRF envelopes that resulted from averaging across the ten split TRFs for each subject. In contrast, Figs. 4, 5, 6 and 7b provide the TRFs and corresponding envelopes for assorted subjects, which were calculated by taking all data of one subject and not the average of the split TRFs.

**Figure 4:**
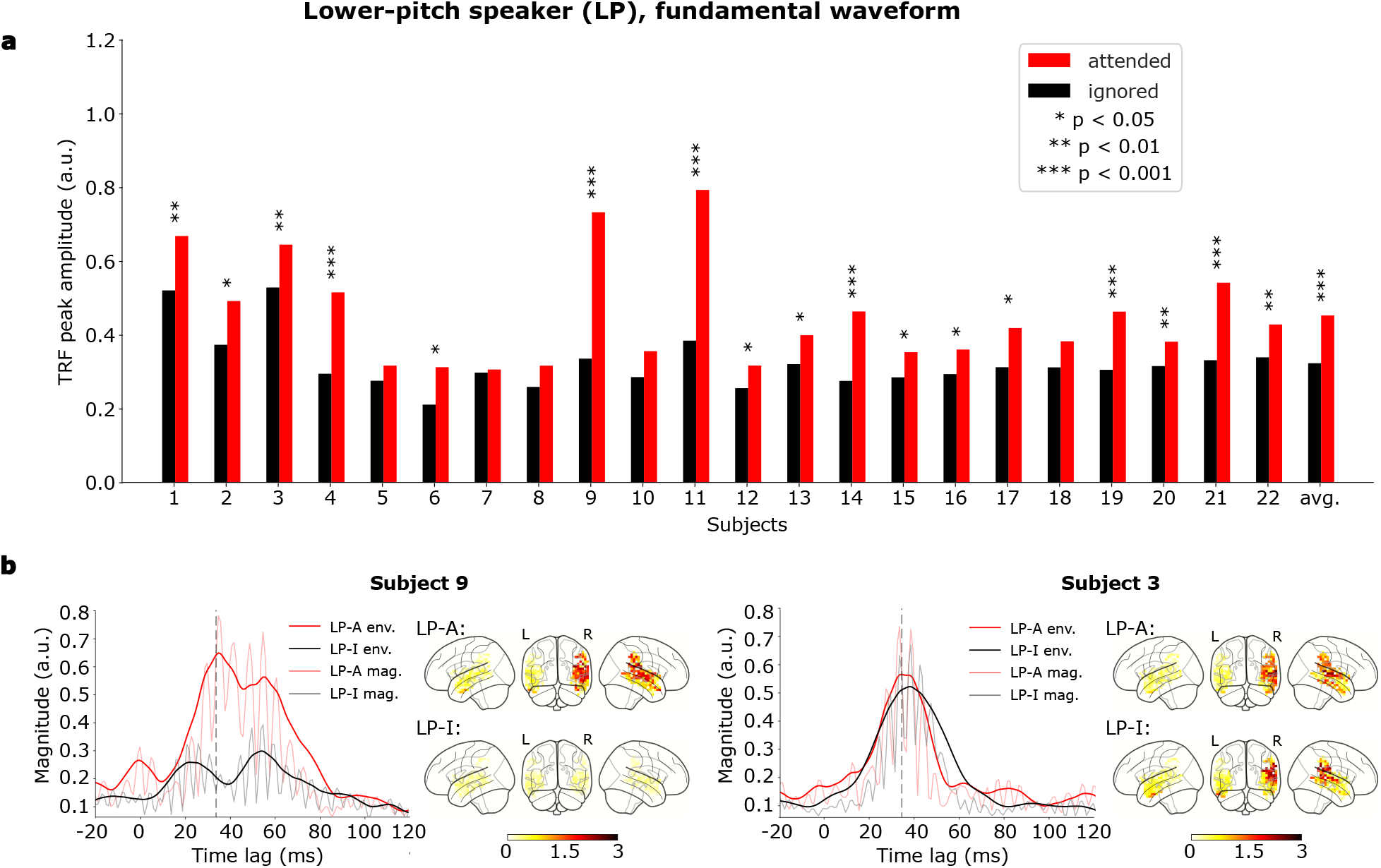
Responses to the fundamental waveform of the LP speaker. (a) Attentional modulation of the cortical contribution to the speech-FFR. For 17 out of 22 subjects the peak in the envelope of the TRF magnitudes at a delay of 34 ms showed a significant difference between the attended (red) and the ignored (black) condition (⋆, 0.01 *≤ p <* 0.05; ⋆⋆, 0.001 *≤ p <* 0.01; ⋆⋆⋆, *p <* 0.001). The same behavior emerged regarding the population-average response (avg.). (b) Cortical TRFs and corresponding voxel magnitudes for the LP-A condition (LP-A env., red; LP-A mag., pink; upper brainplots) and the LP-I condition (LP-I env., black; LP-I mag., grey; lower brainplots) speaker for the exemplary subject 9 (left) and subject 3 (right). There is a large effect of attention for subject 9 and a much smaller one for subject 3.

**Figure 5:**
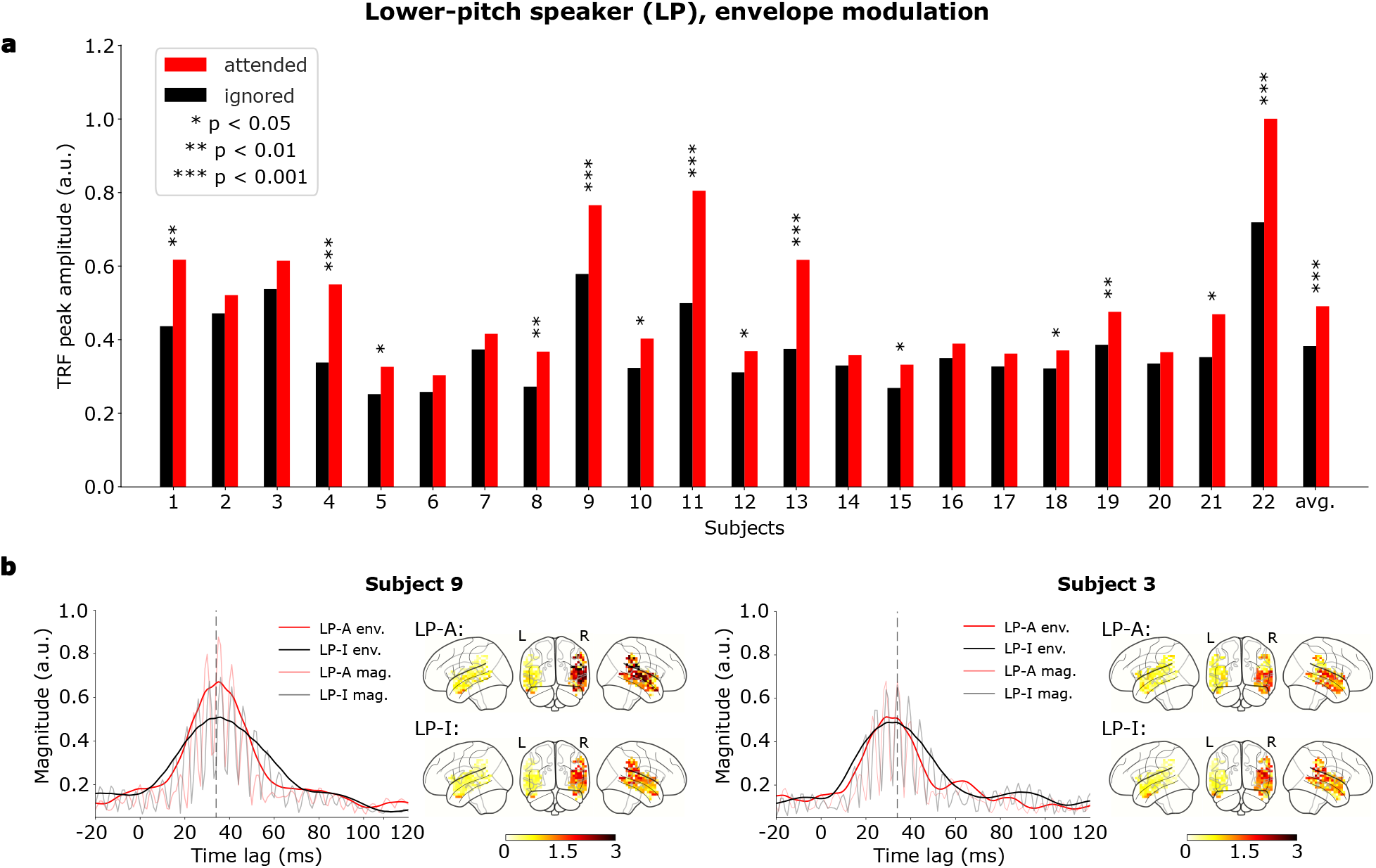
Responses to the envelope modulation for the LP speaker. (a) Attentional modulation of the cortical contribution to the speech-FFR. For 14 out of 22 subjects the envelope of the TRF magnitude at the delay of 34 ms (peak of the envelope) showed a significant difference between the attended condition (red) and the ignored condition (black; ⋆, 0.01 *≤ p <* 0.05; ⋆⋆, 0.001 *≤ p <* 0.01; ⋆ ⋆ ⋆, *p <* 0.001). The population-average TRF (avg.) shows the same attentional modulation. (b) The cortical TRFs and the corresponding voxel magnitudes for the LP-A condition (LP-A env., red; LP-A mag., pink; upper brainplots) and the LP-I condition (LP-I env., black; LP-I mag., grey; lower brainplots) for exemplary subject 9 (left) and subject 3 (right). The channel-averaged TRFs for subject 9 show a strong attentional modulation, while the one for subject 3 is insignificant.

**Figure 6:**
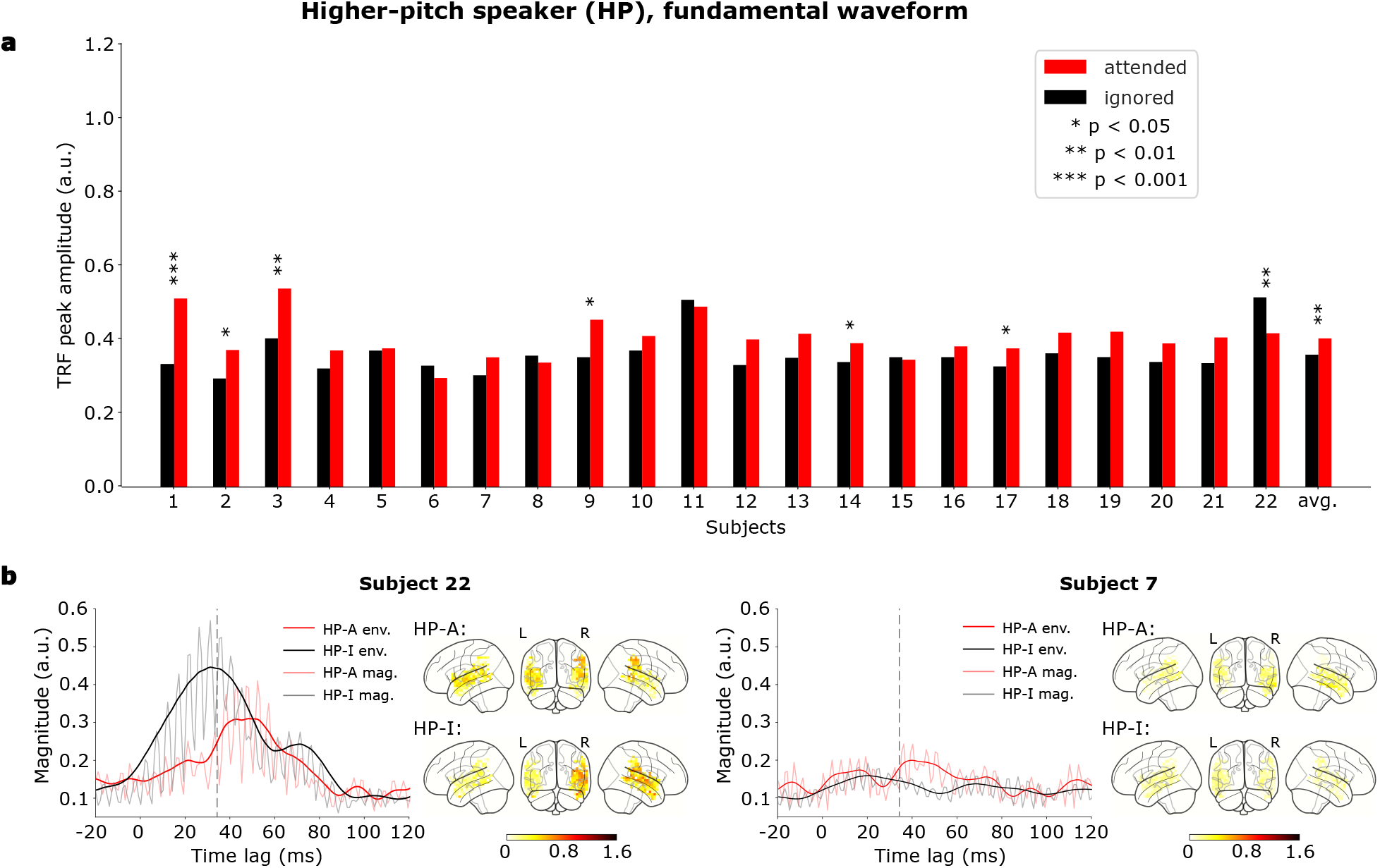
Cortical responses to the fundamental waveform of the HP speaker. (a) Attentional modulation of the cortical response. For 7 out of 22 subjects the peak envelope of the TRF magnitude, at the delay of 34 ms, showed a significant difference between the attended (red) and the ignored (grey) condition (⋆, 0.01 *≤ p <* 0.05; ⋆⋆, 0.001 *≤ p <* 0.01; ⋆ ⋆ ⋆, *p <* 0.001). The population-average TRF (avg.) displayed the attentional effect as well. (b) The time course of the TRF magnitudes and the corresponding envelopes as well as the corresponding voxel magnitudes for the HP-A condition (HP-A env., red; HP-A mag., pink; upper brainplot) and the HP-I condition (HP-I env., black; HP-I mag., grey; lower brainplots) for exemplary subject 22 (left) and subject 7 (right). The cortical response of subject 22 showed a significant effect of selective attention, but not the response of subject 7.

**Figure 7:**
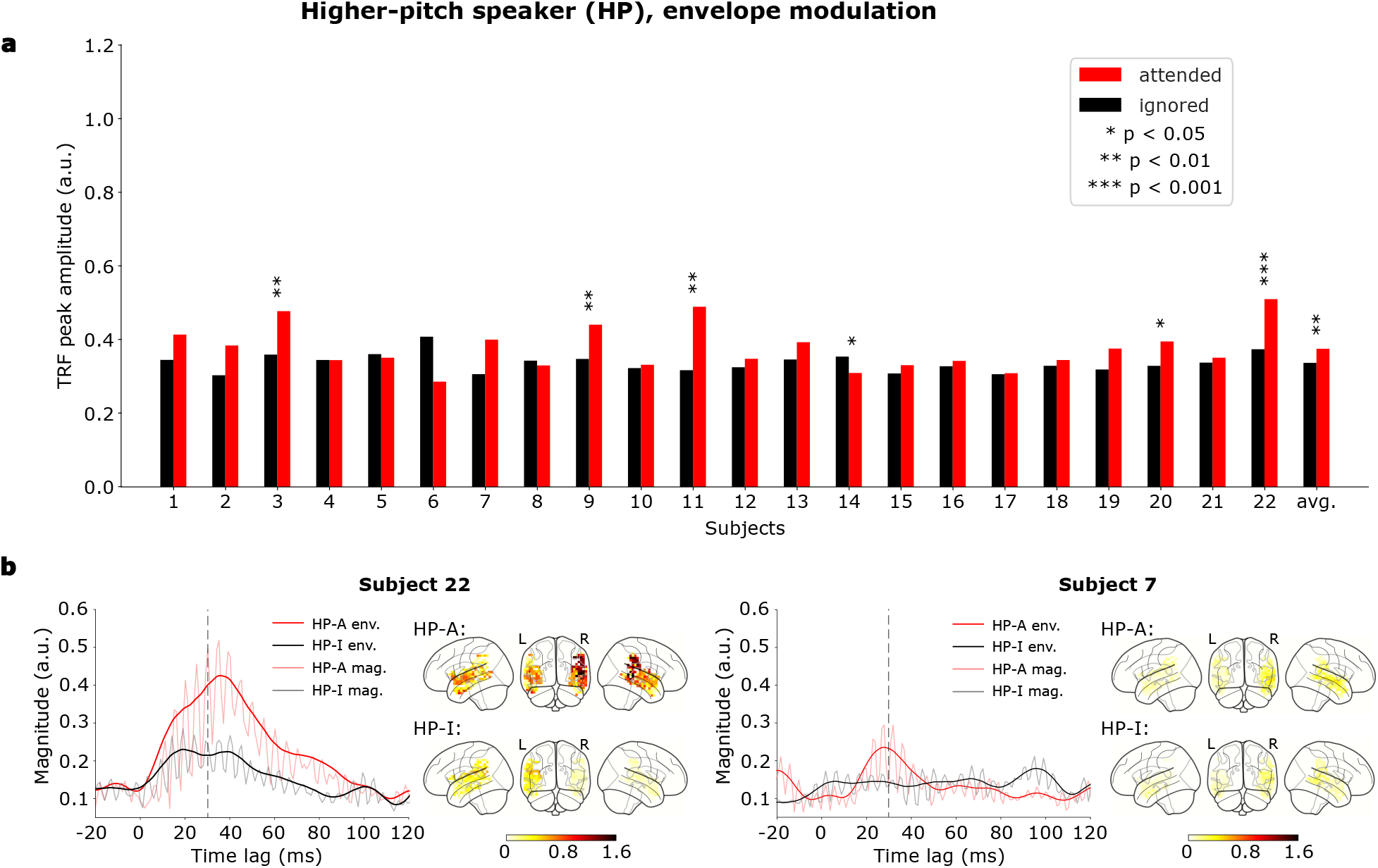
Cortical responses to the envelope modulation of the HP speaker. (a) Attentional modulation of the cortical contribution to the speech-FFR. For 6 out of 22 subjects the peak envelope of the TRF magnitude, at the delay of 30 ms, differed significantly between the attended (red) condition and the ignored (black) condition (⋆, 0.01 *≤ p <* 0.05; ⋆⋆, 0.001 *≤ p <* 0.01; ⋆ ⋆ ⋆, *p <* 0.001). The population average (avg.) showed the attentional modulation as well. (b) Envelopes of the TRF magnitudes as well as the TRF magnitudes themselves and the corresponding voxel magnitudes for the HP-A condition (HP-A env., red; HP-A mag., pink; upper brainplot) and the HP-I condition (HP-I env., black; HP-I mag., grey; lower brainplot) for exemplary subjects 22 (left) and subject 7 (right). The cortical response of subject 22 showed a significant attentional effect, whereas that of subject 7 did not.

Regarding the responses to the fundamental waveform of the LP voice, we found that for most of the subjects (17 out of 22) the envelope of the TRF magnitude at the latency time of interest, 34 ms, when the speaker was attended, significantly exceeded the envelope when the speaker was ignored (Fig. 4a, two-tailed Whitney-Mann rank test). On the population level, the envelope of the TRF magnitudes in the LP-A condition was also significantly higher than in the LP-I condition (*p <* 0.001, two-tailed Wilcoxon signed rank test; Fig. 2, upper left). In particular, the neural response in the attended condition was 26 *±* 3% higher than in the ignored condition (mean and standard error of the mean).

To investigate the time course of the envelope of the TRF magnitudes for individual subjects, as well as the location of the neural sources in the brain, we display this information for two exemplary subjects (Fig. 4b). Subject 9 exhibits significant attentional modulation at the time lag of 34 ms as well as at earlier and later time lags. The neural activity appears predominantly in the primary auditory cortex, and is much smaller when the speaker is ignored. Subject 3 shows a much smaller difference in the neural responses between the attended and the ignored condition.

The subject-specific neural responses to the envelope modulation, at the delay of 34 ms, revealed a similar behavior, with 14 out of 22 subjects displaying a larger neural response in the LP-A condition than in the LP-I condition. The population-average neural response exhibited this behavior as well (Fig. 5a, *p <* 0.001, two-tailed Wilcoxon signed rank test). It was 20 *±* 2% higher in the attended than in the ignored condition (mean and standard error of the mean).

Figure 5b presents further data on the neural response for the same two exemplary subjects as in Fig. 4b. Subject 9 showed again a significant difference between the LP-A and the LP-I condition, whereas subject 3 did not. Compared to the neural response to the fundamental waveform (Fig. 4b), subject 9 exhibited a less noisy signal, especially in the LP-I condition.

Regarding the population-average neural responses to the LP speaker (Fig. 2, upper right), in both the LP-A condition and in the LP-I condition the response to the envelope modulation showed less noise and a higher magnitude and voxel activation than the response to the fundamental waveform (Fig. 2, upper left).

### Attentional modulation of the cortical contribution to the speech-FFR: response to the HP speaker

We applied the same analysis to investigate the attentional modulation of the neural response to the HP voice. We found that, for the response to the fundamental waveform, only a minority of subjects displayed significant attentional modulation (7 out of 22, Fig. 6a). For six subjects, the response in the HP-A condition significantly exceeded the one in the HP-I condition. For subject 22, however, the response in the HP-I condition was significantly larger than in the HP-A condition (Fig. 6b), although both TRFs were noisy. Exemplary subject 7 yielded noisy TRFs both in the HP-A and in the HP-I condition with no significant attentional modulation (Fig. 6b).

The population average (Fig. 2, Fig. 6a avg.) displayed a significant cortical response at the delay of 34 ms as well as a significant effect of attention (*p <* 0.01, two-tailed Wilcoxon signed rank test). The response in the attended condition exceeded the one in the ignored condition by 10 *±* 3% (mean and standard error of the mean).

The neural responses to the envelope modulation, at the peak delay of 30 ms, revealed a similar pattern. Among the 22 subjects, 6 exhibited a significant difference when comparing cortical responses in the HP-A condition to those in the HP-I condition.

The population average of the cortical response displayed a clear peak centered at 30 ms in the HP-A condition, but no clear peak in the HP-I condition (Fig. 2, lower right). It was significantly larger in the HP-A condition than in the HP-I (Fig. 7a, *p <* 0.01, two-tailed Wilcoxon signed-rank test). The difference in the neural response between the two conditions was 8 *±* 4% (mean and standard error of the mean).

Regarding the two exemplary subjects 7 and 22, subject 22 showed a significantly smaller cortical response when the HP speaker was attended as compared to when he was ignored (Fig. 7b). Subject 7, in contrast, did not display a significant effect of attention.

## Discussion

Our study investigated the attentional modulation of the cortical contributions to the speech-FFR, using MEG recordings. We therefore employed a well-established paradigm in which participants focused selectively on one of two competing voices. We then examined the cortical contribution to the speech-FFR through two acoustic features, the fundamental waveform and the envelope modulation. The neural responses were computed through source estimation followed by ridge regression that related the source-reconstructed neural activities to the two speech features. We then assessed the effect of selective attention on these two neural responses.

Our results first verified that both the fundamental waveform and the envelope modulation elicited significant cortical responses. We identified high neural activation in the auditory cortex region for both acoustic features, at latencies ranging from 30 ms to 35 ms. These neural responses were observed despite the presence of a competing speaker, and were presented both for the attended and the ignored voice. These findings are in line with previous research using MEG, where cortical contributions to the speech-FFR in response to short speech tokens were observed (23). In two recent studies focusing on cortical and subcortical contributions to the speech-FFR elicited by continuous speech, cortical responses within similar latency ranges (30 ms to 40 ms) were reported (26; 29).

In contrast to many previous studies on selective attention to competing speakers, we did not employ a male and a female voice, but two male voices. We could thereby demonstrate that selective attention can work also when both competing speakers have the same gender. To differentiate the two voices, we used their pitch, which was higher for one speaker (HP, around 120 Hz) and lower for the other (LP, around 80 Hz). We found that the responses to the LP speaker were larger than those to the HP speaker, indicating that the cortical responses to the speech-FFR decline with increasing fundamental frequency. This finding parallels previous EEG as well as modeling results on the subcortical contribution to the speech-FFR that also showed larger subcortical responses for lower-pitch voices (21; 41; 42). It presumably reflects the decline of phase-locking in the auditory pathway with increasing frequency.

We further found, regarding the response to the LP speaker, that the envelope modulation elicited a larger response than the fundamental waveform. This result emerged both for the attended and the ignored condition, and aligns with a previous finding that the cortical contribution to the speech-FFR is dominated by the response to the envelope modulation (26). Moreover, a recent EEG study found a similar behaviour regarding the subcortical contribution to the speech-FFR (35). Regarding the HP speaker, however, our results showed either a similar response to the envelope modulation and to the fundamental waveform (attended condition) or a higher response to the latter speech feature (ignored condition). This finding could reflect the weaker and thus noisier response to the HP speaker compared to the LP speaker.

The cortical response that we measured was right lateralized, which aligns with prior findings regarding MEG-measured cortical responses to short speech tokens (23) and to continuous speech (26; 43; 29). This right-lateralized pattern may reflect the important role of the right hemisphere in spoken language comprehension, as demonstrated for instance by functional magnetic resonance imaging (fMRI) (44). Additionally, the study by (45) indicated right-lateralized responses in the brain for processing speech and melody, as observed in their fMRI study involving sung stimuli.

Importantly, our results showed a systematic modulation of the cortical contribution to the speech-FFR by selective attention. The clearest results thereby emerged for the responses to the LP speaker, where we observed consistent attentional effects at the peak latency of 34 ms, both for the fundamental waveform and the envelope modulation and on the level of individual subjects as well as on the population level. The LP-A condition led to larger cortical responses than the LP-I condition in all subjects and for both acoustic features, and significant differences between the LP-A and the LP-I condition were observed in more than half of the subjects (Fig. 4a and 5a). Additionally, the neural responses to the fundamental waveform and to the envelope modulation peaked at the same latency, a behavior that was also found on the level of individual subjects (Fig. 4b and 5b). The population average analysis confirmed attentional modulation on a group level for both acoustic features, with the LP-A condition leading to significantly larger responses than the LP-I condition.

Attentional effects on the neural responses to the HP voice were less prominent than for the LP speaker. While we found significant differences between the HP-A condition and the HP-I condition, these effects were not entirely consistent across subjects. For both acoustic features, less than half of the subjects showed a significant difference between the HP-A condition and the HP-I condition (Fig. 6a and 7a). The subject-level TRFs were noisier than those for the LP speaker. In four subjects we found larger cortical responses in the HP-I condition than in the HP-A condition. Importantly, however, the population averages nonetheless yielded a significantly larger response in the attended as compared to the ignored condition.

The consistent attentional modulation that we found on the population level, as well as, in many instances, on the level of individual subjects, aligns with our previous findings regarding attentional modulation of the subcortical contribution to the speech-FFR (19; 20; 21). Moreover, the differences between the neural responses in the attended and in the ignored condition were comparable to those observed on the subcortical level. It hence remains an open question whether the attentional modulation of the cortical contribution to the speech-FFR is entirely driven by the subcortical contribution, or whether further attentional processes act on the cortical response. Recording of the speech-FFR through combined EEG and MEG may in the future be able to clarify this issue through simultaneous high-fidelity measurements of the subcortical and the cortical portion of the speech-FFR.

We would like to highlight that the experimental conditions in our study were highly controlled, with participants switching their attention between two specific speakers, in a supervised environment. However, in real-life situations, attention is a highly dynamic and multifaceted process influenced by various factors such as environmental distractions, cognitive load, and individual differences. In addition, attentional states are often influenced by subjective experiences, cognitive processes, and behavioral cues that may not be directly observable from neural activity. Integrating multimodal approaches such as recently achieved will in the future provide a more comprehensive understanding of attentional processes (46). It will thereby be of particular interest to investigate whether the early cortical response described here will be affected by such additional factors, or if the latter will rather affect cortical processing at a later stage.

## Acknowledgements

This work was funded by the Deutsche Forschungsgemeinschaft (DFG, German Research Foundation): grant KR 5148/2-1 to PK (project number 436456810) and grant SCHI 1482/3-1 (project number 451810794) to AS, and by the Emerging Talents Initiative (ETI) of the University Erlangen-Nuremberg (grant 2019/2-Phil-01 to PK).

## Author contributions

**Alina Schüller**: Conceptualization, Data Curation, Interpretation of Results of Experiments, Formal analysis, Investigation, Writing original draft. **Achim Schilling**: Conceptualization, Interpretation of Results of Experiments, Reviewing and Editing of the paper. **Patrick Krauss**: Interpretation of Results of Experiments, Reviewing and Editing of the paper. **Stefan Rampp**: Interpretation of Results of Experiments, Reviewing and Editing of the paper. **Tobias Reichenbach**: Conceptualization, Interpretation of Results of Experiments, Investigation, Supervision, Reviewing and Editing of the paper.

